# Complementary approaches to dissect late leaf rust resistance in an interspecific raspberry population

**DOI:** 10.1101/2024.01.04.574234

**Authors:** Melina Prado, Allison Vieira da Silva, Gabriela Romêro Campos, Karina Lima Reis Borges, Rafael Massahiro Yassue, Gustavo Husein, Felix Frederik Akens, Marcel Bellato Sposito, Lilian Amorim, Pariya Behrouzi, Daniela Bustos-Korts, Roberto Fritsche-Neto

## Abstract

Over the last ten years, global raspberry production has increased by 47.89%, based on the red species (*Rubus idaeus*). However, the black raspberry species (*Rubus occidentalis*), although less consumed, is resistant to one of the most important diseases for the crop, the late leaf rust caused by *Acculeastrum americanum* fungus, to which the red ones are susceptible. In this context, genetic resistance is the most sustainable way to control the disease, mainly because there are no registered fungicides for late leaf rust in the crop in Brazil. Therefore, the aim was to understand the genetic architecture that controls resistance to late rust in raspberries. For that, we used an interspecific diversity panel between the cited above species, two different statistical approaches to associate the phenotypes to the markers (GWAS and copula graphical models), and two phenotyping methodologies from the first to the seventeenth day after inoculation (high-throughput phenotyping with a multispectral camera and traditional phenotyping by disease severity scores). Our findings indicate that a locus of higher effect possibly controls the resistance to late leaf rust, as both GWAS and the network suggested the same marker. Furthermore, a candidate defense-related gene cluster is close to this marker. Finally, the best stage to evaluate for disease severity is thirteen days after inoculation, confirmed by both traditional and high-throughput phenotyping. Although the network and GWAS indicated the same higher effect genomic region, the network identified other different regions complementing the genetic control comprehension.

## Introduction

Although wild raspberry species occur in different conditions, production is mainly in Europe and North America. With a production increase of 47.89% in ten years (FAOSTAT 2022), this temperate crop is increasingly appreciated due to several characteristics. Its main features are organoleptic properties, content of healthy components, such as anthocyanins (Rao and Snyder 2010), and high added value, which can represent a potential business for small and medium-sized producers (de Oliveira *et al*. 2020). Even with this set of characteristics, there is a need for more raspberry cultivars adapted to tropical climate conditions, as is the case in Brazil. In Brazil, the used varieties are introgressions from north-hemisphere countries (Marchi *et al*. 2019), so their yield and performance in the face of biotic and abiotic stresses depend on the intensity of the interaction of these cultivars with the new production environments. The country’s production is concentrated in regions with high altitudes and lower annual temperatures, representing a small portion of arable areas of Brazil.

The acceptance of a new raspberry cultivar relies on environmental adaptation and fruit quality, with particular emphasis on fruit color. Red raspberries (*Rubus idaeus*) are economically more important and more consumed than black ones (*Rubus occidentalis*) (Graham and Woodhead 2009; Baby *et al*. 2018). However, in addition to the differences in fruit color, they also differ in resistance to some diseases. Late leaf rust is relevant in the north and south hemispheres, with reported losses of up to 70% with the cultivar “Festival” in Nova Scotia (Ellis *et al*. 1991). The *Acculeastrum americanum* pathogen causes the disease, and the first symptom is the orange spots on the leaves, which turn dark over time. Depending on the disease severity, premature defoliation may occur, consequently becoming more susceptible to winter injuries. Although most red raspberry cultivars are susceptible to late rust and are economically important, more resistant varieties, such as the black raspberry species, have been reported and may serve as a source of genetic variability (Hall *et al*. 2009). Therefore, exploring the resistance interspecific variation to late leaf rust in raspberries represents great potential for breeding and a more sustainable and indicated way to control the disease.

The different genetic variations in these species derive from spontaneous mutations maintained or shaped during evolution by some forces, such as natural or artificial selection. The analysis of these mutations helps to elucidate the genetic architecture of traits of great interest to humanity, such as yield or resistance to biotic and abiotic stresses, among many others (Alonso-Blanco *et al*. 2009). Understanding complex traits requires a study of the causal loci allelic variation. The general way to make this link between phenotype and genotype is by performing broadly used analyses, Genome-Wide Association Studies (GWAS), and QTL mapping (Korte and Farlow 2013). The wide use of these techniques was possible due to the advancement of next-generation sequencing technologies, exponentially increasing the capacity to obtain genomic data with a drop in the associated cost (Henson *et al*. 2012). However, phenotyping remains a bottleneck for these traits studied nowadays (Yang *et al*. 2020). Our ability to dissect the genetic architecture of a quantitative trait is limited by our ability to obtain individual phenotypic values, such as traditional or high-throughput phenotypes. As traditional methods can be costly, time-consuming, destructive, and more prone to human error, high-throughput phenotyping has come to try to reduce the phenotyping bottleneck in breeding programs (Gill *et al*. 2022). Although it still presents many limitations and challenges, once established in a crop, it can accelerate genetic gain by improving the selection intensity and accuracy (Araus *et al*. 2018).

In the past years, GWAS has been the main analysis used to study the genetic architecture of traits due to its ability to control population structure and not require a specific mating design as QTL mapping. The main contribution of the technique derives from the fact that it is possible to perform these associations in panels containing great diversity. The method uses the ancestral recombination events to identify the causal loci through the linkage disequilibrium (Huang and Han 2014). The studied traits can be simple and easily detectable by GWAS when they have few loci with large effects on the phenotype or more complex traits, such as those that are regulated by many small-effect loci or those that have many rare allelic variants (Korte and Farlow 2013).

Not as conventional as associative and QTL mapping in plant breeding, it is the use of a graphical network. The term *Network* has become more frequent in many scientific fields because graphical modeling is one of the best ways to represent high-dimensional data (Purutçuoğlu and Farnoudkia 2017). Behrouzi and Wit (2019) studied epistatic selection using semiparametric penalized copula Gaussian graphical models, in which the network was able to capture aberrant associations between markers through shortand long-range linkage disequilibrium dependencies. Zhang *et al*. (2021), also using latent Gaussian copula models, studied schizophrenia using genomic data based on Single Nucleotide Polymorphism (SNPs) and brain imaging. In these systems, each variable, such as phenotype and genotype, is represented by a node, and edges between these nodes represent its conditional independence relationships. According to Sklar’s theorem, it is necessary to describe the joint probability distribution function to use multivariate stochastic models. This joint distribution can be decomposed into the variable’s marginal distributions and a joint behavior of the random variables, the copula (Durante *et al*. 2013). Besides the capability to utilize data with missing values, a key benefit lies in the separate and independent modeling of marginal distributions for each variable. This approach treats each variable as having a distinct distribution, leading to individualized modeling for enhanced accuracy. As associative mapping and the network derived from copula graphical models are approaches that differ in methodology and data modeling, using them together can contribute as an additional layer of reliability to the obtained results.

Given the above, this work seeks to find the genomic regions responsible for resistance to late leaf rust in raspberries using traditional and high-throughput phenotyping data through a nonconventional pipeline using a genome-wide association study and copula graphical models network.

## Materials and methods

### Plant material

The crosses to obtain the interspecific hybrids were carried out in a *testcross* scheme, in which a group consisted of 3 parents of the species *R. idaeus* that had favorable characteristics for the market, such as fruit color ranging from golden (“Golden Bliss”) to red (“Himbo Top”), were crossed with the central parent of the species *R. occidentalis* (“Jewel”), being the source of alleles for late rust resistance (Figure 1). The number of crosses obtained with each parent of the first group was varied. “Jewel” was used as a female parent due to the unilateral incompatibility between the species (LEWIS and CROWE 1957). Of the 99 genotypes, 94 were interspecific hybrids, and 4 were parents. More information about the genetic characterization and diversity of the panel can be found in Campos *et al*. (2023).

**Figure 1.**
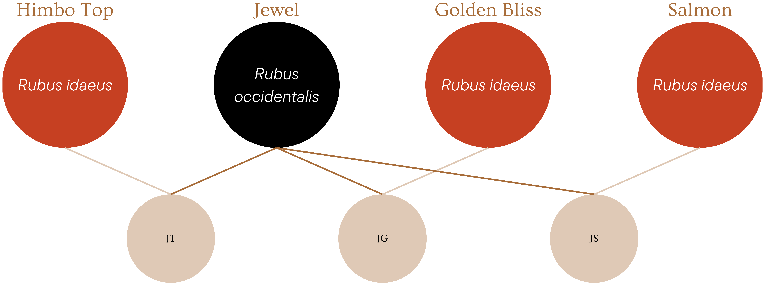
Population structure. In red are the varieties of *Rubus idaeus*, with different susceptibility levels. In black is the central parent of the species *Rubus occidentalis*, mother of all crosses, with higher resistance to late-leaf rust.

The variety “Jewel” is described as a vigorous, consistently productive plant with large, firm fruits, almost black color, and greater resistance to winter injuries than red raspberries (OURECKY; SLATE, 1973; OMAFRA, 2021). The species *R. occidentalis* has late rust resistance records, but the records are not associated with the “Jewel” variety. The “Heritage” variety has medium, reddish, and excellent quality fruits, but it is highly susceptible to late rust and needs more hours of chilling (Hall *et al*. 2009). The cultivar “Himbo Top,” which originated in Switzerland, has pale red fruit and is easy to harvest (OMAFRA, 2021; HAUENSTEIN, 2008). The cultivar “Autumn Bliss,” parent of “Himbo Top,” has good productivity in Brazil and is less demanding in terms of the number of chilling hours than “Heritage” (Raseira *et al*. 2004). The “Golden Bliss” variety, or” All Gold” in the United Kingdom, has little information in the literature. It has yellow fruit, and its possible parent is the cultivar “Autumn Bliss” (GIRICHEV, 2017). The variety was well adapted to the South of the state of Minas Gerais (MOURA et al., 2012). No information about the “Salmon” variety was found in the literature.

### Conducting the experiments

The experimental units consisted of a 5-liter pot containing only one plant. The experiment was arranged in an augmented block design, in which the parents were replicated several times and in a systematic way. The experiment was carried out in a semi-controlled condition, in a greenhouse with temperature control and supplemental light. The checks, present in each of the three blocks, consisted of a clone of the “Golden Bliss” variety, one of the “Himbo top,” one of the “Heritage,” one of the “Jewel,” and one of the “Salmon” variety. The experiment was replicated twice in time. For its execution, drastic pruning was carried out, leaving only three buds on the stems. Because younger leaves are less susceptible to late rust, with the drastic pruning, all the plants had leaves at approximately the same development stage. The inoculum from the fungus *Aculeastrum americanum* was sprayed five days after the pruning. The suspensions were prepared in the Department of Phytopathology of ESALQ/USP, using 50 ml of distilled water, Tween 20 (0.01%), and urediniospores of *A. americanum* collected. The suspension concentration was adjusted to 104 urediniospores/mL in a Neubauer chamber and used to spray inoculate the abaxial face of the leaves to the point of runoff. To ensure the development of the disease, the plants were covered for 24 hours with a dark plastic bag to set up a humid chamber.

### Phenotyping

The experiment consisted of two types of phenotyping, traditional and high-throughput phenotyping, lasting approximately 17 Days After Inoculation (D.A.I.) (Figure 2). In traditional phenotyping, disease severity scores (ranging from 0 to 8) were assigned from the eleventh to the seventeenth D.A.I. According to the diagrammatic scale proposed by (Dias *et al*. 2023a). Since there is subjectivity related to the score given by the evaluator and to reduce this bias, three people evaluated all the plants and an average of the evaluators was performed.

**Figure 2.**
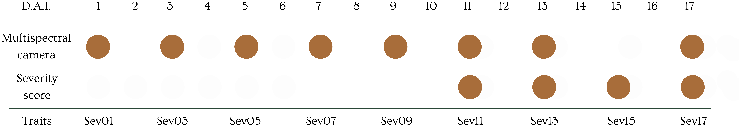
Replicate structure. The experiment was replicated twice, and there were 17 days of phenotyping after inoculation in each replicate. We phenotyped in two ways: traditional phenotyping using disease severity scores and high-throughput phenotyping using the multispectral camera.

A high-throughput greenhouse phenotyping platform obtained the plant’s multispectral images (Figure 3). The platform consists of 3 rails, two moving in parallel and one perpendicular to the side of the greenhouse. A board, cameras, battery, and sensors were attached to the perpendicular rail. More details about this low-cost, greenhouse-based, high-throughput phenotyping platform in Yassue *et al*. (2021).

**Figure 3.**
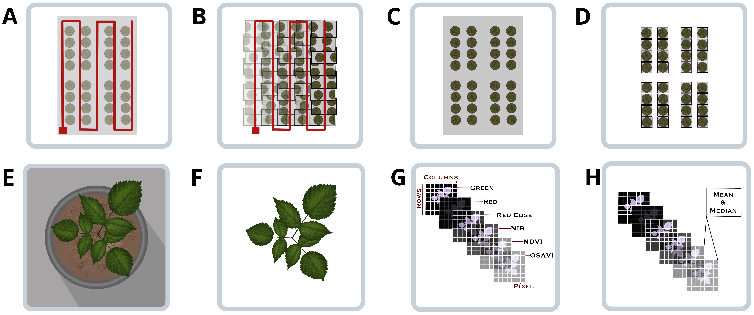
High-throughput phenotyping methodology. A - Mimicry of a drone flight inside the greenhouse using motorized rails and the multispectral camera. B - Georeferenced images with 70% overlapping. C - The orthomosaic assembly. D - The extraction of ShapeFiles from orthomosaic (pots). E - The image of a plot with the background. F - The image of a plot after removing the background. G - This same image is represented with a 3D matrix, where the third dimension represents the original and calculated layers (masks). The spectral indices (phenotypes) are calculated through the mean and median of these layers.

The high-throughput images were collected from the first Day After Inoculation (D.A.I.) and then on alternate days until completing 17 D.A.I. We needed help to retrieve data for one of the high-throughput phenotyping days, the fifteenth D.A.I. To facilitate discussion in the paper, we will design these traits as Sev01 to Sev17, specifying the first to the seventeenth phenotyping D.A.I. for disease severity.

### Image processing

The multispectral camera was the *Parrot’s Sequoia+*, with a resolution of 1.2 MP in each of the four monochromatic lenses: green, red, red edge, and near-infrared. The monochrome lens of the green has a wavelength of 550 nm, the red has a wavelength of 660 nm, the red edge of 735 nm, and the near-infrared of 790 nm. The images were processed by assembling the orthomosaics using the *software AgiSoft Metashape v. 1*.*7*. After assembly, the plots *ShapeFiles* were drawn in the *software QGIS 3*.*0*. To remove the background, we first apply the *Soil Color Index* (SCI) filter for soil removal with the software *FieldImageR* (Matias *et al*. 2020). As the image’s background was uneven, we had to manually adjust the intervals of each layer to remove as much of the background as possible. The spectral indices referring to the pre-symptomatic and symptomatic plants were extracted using the same software. We tested two different masks, the NDVI and OSAVI, from which we extracted the two indices commonly used in works studying disease severity. We used the mask layers’ mean and median, as carried out by Yassue *et al*. (2021) and as represented in Figure 3 (step H).

### Genotyping

Genotyping was performed using the *Genotyping-By-Sequencing* methodology (GBS) (Elshire *et al*. 2011). Young fresh leaves were sampled from each genotype and immediately frozen in liquid nitrogen. These samples were stored at −80°*C* until further analysis. The extraction of total DNA from each sample was carried out following the protocol proposed by the manufacturer of the extraction kit (*Qiagen*) and using the *DNeasy Plant Mini Kit*. A rare-cutting enzyme, the PstI (New England BioLabs Inc.®), and another frequently-cutting enzyme, the MseI (New England BioLabs Inc.®), were used. Purification with the *QIAquick PCR purification Kit* (*Qiagen*) was followed to proceed with the PCR amplification step. DNA libraries were quantified using the *Agilent DNA 1000 Kit* in an *Agilent 2100 Bioanalyzer* (Agilent Technologies). Additionally, they were quantified on a *CFX 384 Touch Real-Time PCR Detection System* using the *KAPA Library Quantification* Kit (*KAPA Biosystems, cat. KK4824*). Finally, the libraries were diluted and sequenced in a sequencer *HiSeq 2500 System* (*Illumina, Inc*). The SNP calling was carried out following the pipeline *TASSEL-GBS* (Glaubitz *et al*. 2014). The alignment was performed using *Bowtie2 software* (Langmead and Salzberg 2012) with the reference genomes of the black raspberry (VanBuren *et al*. 2016). Filters, such as minor allele frequency ≥ 0.20, missing rate ≤ 0.20, and a filter to remove monomorphic SNPs, were applied to remove possible sequencing errors using the package *SNPRelate* (Zheng *et al*. 2023). We used this MAF value because it helped to control the large variability in the panel and improve the resolution of the GWAS in our small population and because we were interested in the resistance alleles of the central parent. We used 28.373 markers from the original data (Campos *et al*. 2023) and ended with 19.440 markers after filtering. Finally, the snpReady software was used for the missing data imputation by Wright’s method (Granato *et al*. 2018).

### Phenotypic analysis

The genotypic values and variance components were estimated using the REML/BLUP method (*Restricted Maximum Likelihood/ Best Linear Unbiased Predictor*) using the *statgenSTA* package (Rossum *et al*. 2023) in the R environment (version 4.3, https://www.rproject.org/). The following model was used to calculate the genotypes BLUPs, to estimate the variance components and Cullis heritabilities:

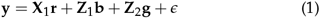

Where **y** is the vector of phenotypic values of disease severity; **r** is the replicate fixed effect; **b** is the random replicate nested block effect, where 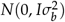; **g** is the random genotype effect, where 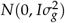; *ϵ* is the random effect of the residual, where 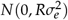, **X**_**1**_ is the fixed effect incidence matrix; **Z**_**1**_ and **Z**_**2**_ are the random effect incidence matrices; **I** is the identity matrix; **R** is the variance and covariance matrix of the residual effects. BLUPs were considered in the next steps because it was necessary to perform a spatial correction using checks with the *statgenSTA* package, due to the heterogeneity of spatial effects and the experimental design in the greenhouse. As *statgenSTA* package uses *SpATS* package as a modelling engine and we modelled the spatial effects, a P-spline ANOVA (PSANOVA) approach function from the last package was used (Rodríguez-Álvarez *et al*. 2017).

### Copula Graphical Model

The *netgwas* package was used to estimate the conditional independence relationships with a nonparanormal (“npn”) approach within the Gaussian copula graphical model (Behrouzi *et al*. 2023), this method was chosen because of the data high dimensionality. We built ten networks with different sparsity levels from the following “rho” settings: 0.1, 0.2, 0.25, 0.3, 0.4, 0.5, 0.6, 0.65, 0.7, 0.75. After building the ten networks, we used a package function that calculates the best regularization parameter (rho) based on Bayesian criteria, which was 0.2 for our data. As input for the network, we used a genotype by variable matrix, with the variables being all the BLUPs from the model (1) and all markers. Since copula graphical models are computationally intensive, we reduced the variables number. The *SNPRelate* package was used to filter the marker matrix by Linkage Disequilibrium (LD), using a sliding window with a size of 100 kb and an LD threshold of 0.10, resulting in a 4686 markers matrix (Zheng *et al*. 2023). We ensured that the SNPs found by GWAS were in this new marker matrix, so we could compare the results. To facilitate the discussion of the resulting network, we adopted a partial correlation threshold of 0.05 between the markers and the traditional phenotypes (Sev11 to Sev17). Finally, we used the *netShiny* package to graphically visualize the network (de Jongh and Behrouzi 2022).

### Genome Wide Association Studies

From model (1), associative mapping was performed using the Bayesian information and Linkage disequilibrium Iteratively Nested Keyway (BLINK) method, implemented in the *GAPIT software* (Lipka *et al*. 2012). Quantile-quantile and Manhattan plots were generated to scan the population stratification and visualize the significant SNPs in the associative mapping analysis. We calculated a significance threshold for each trait using the “Farm-CPU.P.Threshold” function (Liu *et al*. 2016). We did not use any principal components as covariates in the model.

### Gene annotation

All candidate genes flanking the associated marker were analyzed considering a 100 kb window, or 50 kb upstream and downstream regions of the physical positions of the associated SNPs. These genes were annotated by using the *R. occidentalis* reference genome. In addition to gene annotation by homology using BLAST, the conserved domains/motifs of R genes from the published *R. occidentalis* genome were predicted by the *InterProScan v5*.*33–72*.*0 software*. According to the literature, candidate genes were classified as not defense-related, belonging to the signaling cascade, as receptors or defense executors.

### SNPs heritability

To estimate the percentage that all associated SNPs explain of the phenotype, we have calculated the SNPs variance components based on the following model:

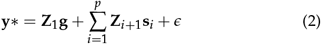

To estimate the percentage that each associated SNP explains of the phenotype, we have calculated the SNPs variance components based on the following model:

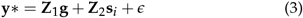

Where **y**∗ is the vector of the disease severity BLUPs calculated in model (1); **g** is the random genotype effect, where 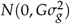 and *G* is the additive kinship matrix; **s**_*i*_ is the random effect of the ith SNP, where 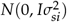; *ϵ* is the random effect of the residual, where 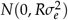; **Z**_1_, **Z**_*i*+1_, …, **Z**_*p*+1_ are the random effect incidence matrices; **I** is the identity matrix; **R** is the variance and covariance matrix of the residual effects.

From model (2), we calculated the narrow sense heritability for all SNPs jointly by the following model:

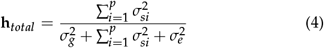

From model (3), we calculated the narrow sense heritability for each SNP:

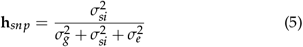

## Results

### Traditional traits

#### Severity score BLUPs

The increase in mean disease severity score BLUPs shows that we were able to capture disease progression with the traditional phenotyping method used (Figure 4). For the same BLUPs, the lowest correlation found was between Sev11 and Sev17, with a correlation of 0.88, and the highest correlation was 0.97 between the Sev15 and Sev17 traits. Cullis heritabilities ranged from 0.36 to 0.40, these values being for Sev17 and Sev13, respectively (Table 1).

**Table 1.** Pearson correlation and Cullis heritability of severity score BLUPs from the eleventh to the fourteenth day after inoculation.

|  | Traits |  |  |  | Cullis Heritability |
| --- | --- | --- | --- | --- | --- |
|  | Sev11 | Sev13 | Sev15 | Sev17 |  |
| Sev11 | 1 |  |  |  | 0.38 |
| Sev13 | 0.96 | 1 |  |  | 0.40 |
| Sev15 | 0.92 | 0.96 | 1 |  | 0.39 |
| Sev17 | 0.88 | 0.93 | 0.97 | 1 | 0.36 |

**Figure 4.**
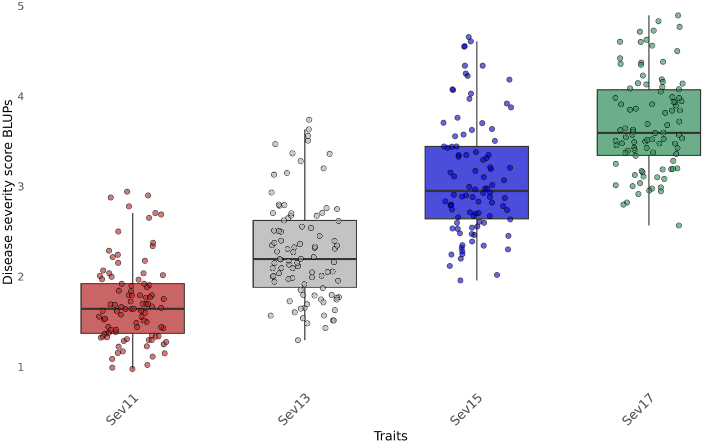
Severity score BLUPs distribution from the eleventh to the seventeenth day after inoculation, or for Sev11 to Sev17 traits.

#### Genome-wide association studies for traditional traits

We found a significant marker with 13 D.A.I., this marker also appears with the highest p-value in all traditional phenotyping days (Figure 5). We already expected to find fewer SNPs because it is a small population, so we decided to monitor this SNP because it is consistent over time. To facilitate the results presentation in this section, we will comment on genes directly related to plant immunity. However, in the next sections, we will demonstrate the gene profile of all genes found by GWAS and the network 8). A list of the entire annotation can be found in the supplementary material for all genes flanking the marker region in a 100 kb window (Table S1). This significant marker is at position 13.3 Mb of chromosome 5. Of the 12 genes flanking its region, 4 were possible receptors, three were likely defense executors, one gene was part of signaling cascades, and four were classified as non-defense related. Approximately 20kb from this marker is a gene whose function prediction is “NB-ARC DOMAIN DISEASE RESISTANCE PROTEIN.” In the exact position of the marker, there is another gene with the function “PLANT BROAD-SPECTRUM MILDEW RESISTANCE PROTEIN RPW8”. 20 kb away, we found a receptor gene with the function “L-TYPE LECTIN-DOMAIN CONTAINING RECEPTOR KINASE S.5-RELATED”. 6.8kb away from the marker, we found the fourth gene with receptor function, the gene “WALL-ASSOCIATED RECEPTOR KINASE-LIKE 21”. In addition to the Manhattan plots, we observed that the QQPlot in 13 D.A.I. is adjusted (Figure S1), with the majority of markers having an expected p-value and only one marker with the observed p-value much higher than expected, supporting the significance of this marker on chromosome 5.

**Figure 5.**
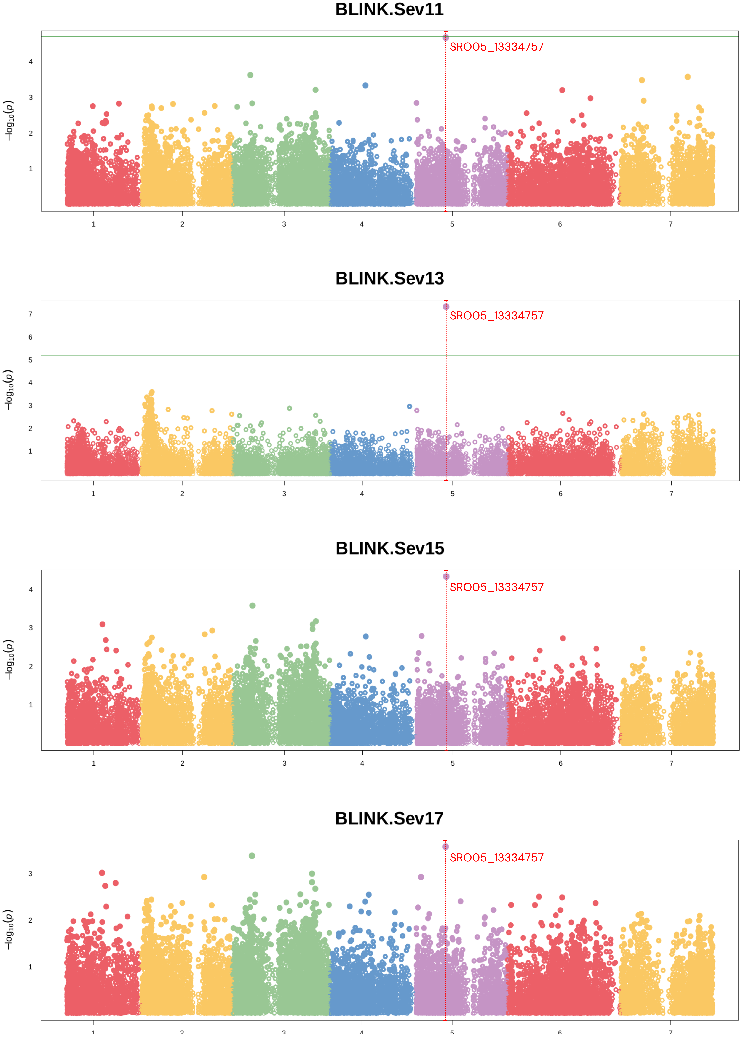
Manhattan Plots from the Genome-Wide Association Studies for each of the four traditional traits, Sev11 to Sev17. The vertical red dotted line represents the marker with the highest p-value. The horizontal green line is the significance threshold calculated for each individual trait.

#### Copula graphical models network

The network resulting from the copula graphical models is in Figure 6. The nodes represent all the variables, the four traditional traits, and the 4686 markers used, while the edge thickness represents the degree of partial correlation between variables. From left to right, we show the network containing all the variables in the A subfigure. The B subfigure shows a subnet containing only the markers with primary dependencies on the four traditional traits. Finally, we show again a subnet in the C subfigure, but having variables with a partial correlation higher than or equal to 0.05. We chose only partial correlations of at least 0.05 to facilitate the discussion of the work. In this same image, the edges connecting the four traditional traits are thicker and have a higher correlation than the correlations with the markers. As with GWAS, annotations for all associated markers are presented in the supplementary material (Table S1).

**Figure 6.**
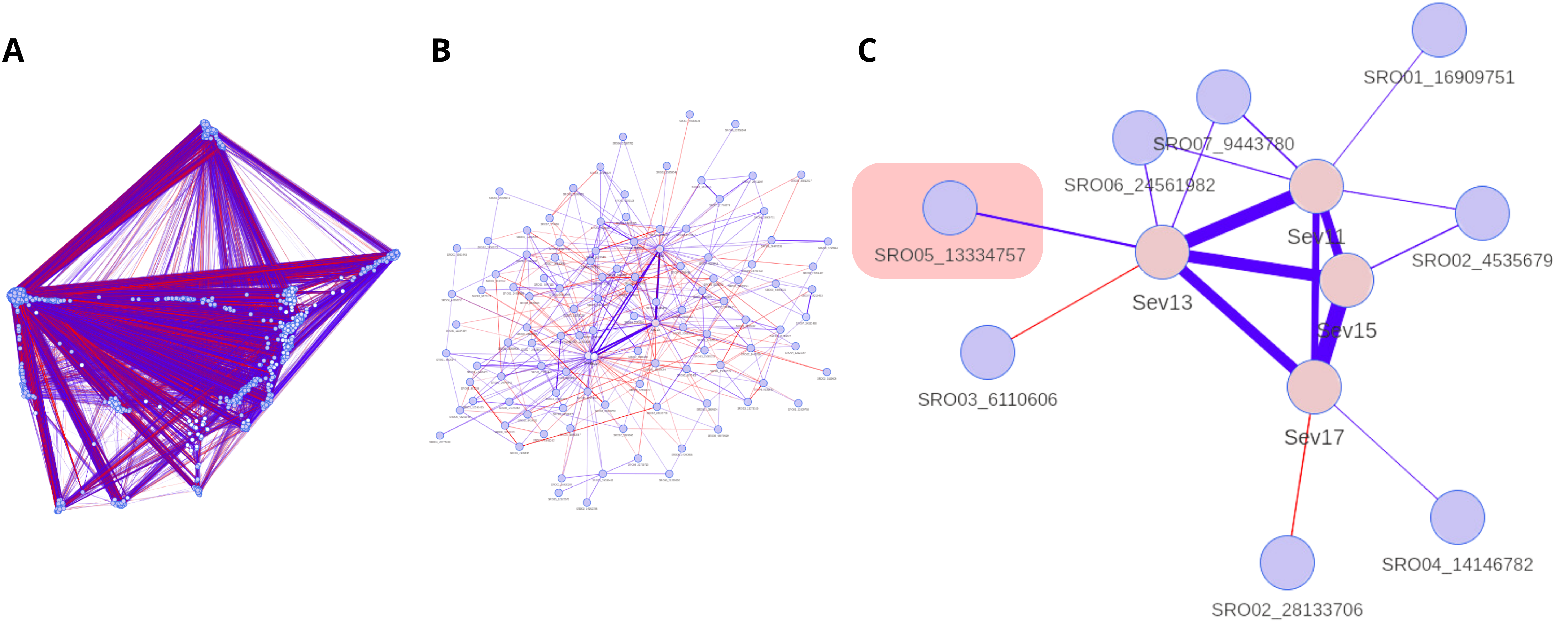
Copula graphical models networks. The A subfigure is the network containing all variables. The B subfigure contains only the markers with primary dependencies on the four traditional traits. The C subfigure is again a subnetwork containing variables with a partial correlation higher than or equal to 0.05. The pink square in the C subfigure highlights the same significant marker in GWAS.

The marker with the higher partial correlation in the network is the SRO05_13334757 marker in 13 D.A.I. (Sev13) (Figure 6 and 7), the same as in GWAS. But unlike GWAS, the network provided us with more complex information about the architecture of late rust resistance. We observed that the four phenotypic variables (In pink in Figure 6, subfigure C) are linked to at least one marker (In blue in Figure 6, subfigure C), and that some phenotypic variables have more than one marker linked to them at the same time. In the partial correlation matrix in Figure 7, some associations are positive (blue) and others negative (red), suggesting that disease severity is negatively or positively correlated with the lack or presence of the reference allele or the most common allele at a given SNP.

**Figure 7.**
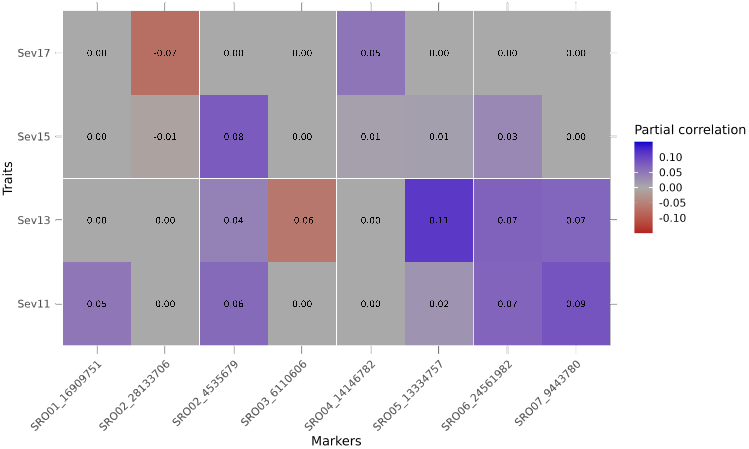
Matrix representation of partial correlations higher than 0.05, between phenotypic and marker variables. The closer the partial correlation is to blue, the more positively related the allele is to the traits. The closer the partial correlation is to red, the more negatively related the allele is to the traits. The gray color represents a partial correlation of zero between the variables.

#### Markers’ gene profile

Figure 8 shows a gene profile for each associated marker, as the regions flanking the eight markers of interest had 93 genes. Of this total, we classified each gene as “Not defenserelated,” Receptors,” Signaling cascades,” and “Defense executors” based on the literature. A table containing our classifications, the classified genes, and the reference articles are available as supplementary material (Table S1). Of the 93 genes, we classified 33 genes as “Not defense-related.” The gene with the highest number of “Defense Executors” in red is the SRO06_24561982 marker. The gene with the highest number of “Receptors” in green is the SRO05_13334757 marker. The gene with the highest number of “Signaling cascades” in blue was the SRO02_4535679 marker.

**Figure 8.**
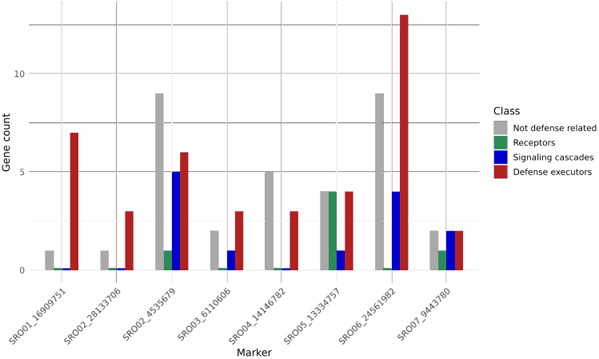
The marker’s gene profile in GWAS and the copula graphical models network. In gray is the not defense-related gene count for each marker on the X-axis. In green is the gene count that has a receptor function. In blue is the gene count that participates in signaling cascades. In red is the gene count with functions belonging to the class of defense executors genes.

#### Graphical network gene annotation

The four phenotypic variables in the network are linked to eight different marker variables, these markers have 93 candidate genes flanking them. The phenotypic variable Sev11 has partial correlations with four different markers (Figure 6, subfigure C). As an example, they have candidate genes with functions such as “ankyrin repeat-containing protein NPR4”, “probable leucine-rich repeat receptor-like protein kinase At1g68400”, “probable WRKY transcription factor 2”, “pleiotropic drug resistance protein 1”, and “salicylic acid-binding protein 2” (Table S1). Just like the Sev11 variable, Sev13 has four markers linked to it, with functions such as “probable serine/threonine-protein kinase kinX” and all the other candidate genes already mentioned for the highlighted marker on chromosome 5. Unlike previous markers, the Sev15 phenotypic variable has only one marker linked to it, with candidate genes such as “probable serine/threonine-protein kinase kinX” gene. Finally, the Sev17 variable has a partial correlation with two markers that are not common to any of the other phenotypic variables and that only have genes that are likely defense executors, as the “FAR-RED IMPAIRED RESPONSIVE (FAR1) FAMILY PROTEIN” gene (Figure 8).

#### SNPs heritabilities

We calculated the heritability for eight SNPs, the marker found in GWAS, and the other 7 SNPs found only by the network. The total for Sev11 was 0.5135, while for Sev13, Sev15, and Sev17, it was 0.5850, 0.5831, and 0.5826, respectively. For SRO05_13334757 marker, the **h**_*snp*_ on the different phenotyping days were 0.3739, 0.4313, 0.3852, and 0.3477. The SRO06_24561982 marker and the chromosome 5 marker obtained the highest heritabilities over the phenotyping days and had heritabilities values of 0.3205, 0.3486, 0.3074, and 0.3013, respectively.

### High-throughput phenotyping

#### Genome-wide association studies for high-throughput traits

Of all the spectral indices used to perform GWAS, only six have a significant marker (Table 3). The heritabilities of these indices ranged from 0.00 to 0.30, while the correlation with traditional traits ranged from 0.08 to 0.23. The only index with a heritability higher than 0.30 were two indices with 13 D.A.I., namely the mean OSAVI, and mean NDVI indices. The same significant marker appears for these two indices: SRO01_14234912 marker is found at position 14 Mb on chromosome 1 (Table 3). Close to this marker, we found a “cysteine-rich receptor-like protein kinase 3”, “auxin response factor 8,” and “Salt stress response/antifungal” genes. We noticed that both NDVI and OSAVI masks produced the same results.

## Discussion

### The polygenic architecture of late rust resistance

We aimed to understand the genetic architecture regulating resistance to late leaf rust in raspberries using traditional and highthroughput phenotyping, using statistical techniques such as GWAS and copula graphical models. The GWAS results showed a higher effect marker on chromosome 5 (Figure 5). Still, there is evidence through copula graphical models that this trait can be regulated, not only by this higher effect marker but also for different genomic regions depending on the stage of disease progression (Figure 7). Studies using other crops in the literature show that rust resistance is polygenic. Ledesma-Ramírez *et al*. (2019) found 14 resistance-associated SNPs to yellow rust in wheat, explaining 6.0 to 14.1% of the resistance. Vikas *et al*. (2022) found 65 quantitative trait nucleotides (QTNs) associated with leaf rust resistance in bread wheat, explaining 1.98–31.72% of the phenotypic variation. We only found one associated marker in GWAS, which explains 43.13% of the phenotypic variation in 13 D.A.I. (Table 2, SRO05_13334757 marker), being a good candidate to carry out marker-assisted selection in raspberry breeding. We believe the small population used is why we did not find other SNPs. Although we knew that a larger population would be ideal, we lost many materials because the experimentation site is located in a subtropical region of Brazil. In contrast, the production sites have a predominantly temperate climate. In addition to the climatic difficulty, the interspecific hybrids’ vigor was unknown as we worked with the F1 population. Even with the reduced population, the significant marker found by GWAS has nine candidate defense-related genes, in addition to other possible regions found by the network (Figure 8). Clustering genes participating in the same metabolic pathway implies they are under the same selective pressure (Polturak and Osbourn 2021). Possibly, not just one of these genes confers resistance to raspberry, but others from the same cluster for a coordinated expression of specialized metabolites (Bharadwaj *et al*. 2021).

**Table 2.**
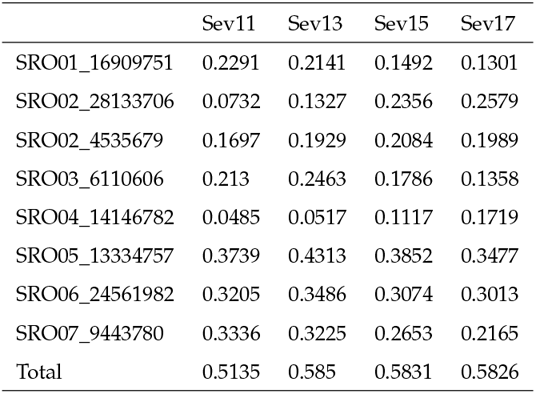
SNPs heritabilities.

|  | Sev11 | Sev13 | Sev15 | Sev17 |
| --- | --- | --- | --- | --- |
| SRO01_16909751 | 0.2291 | 0.2141 | 0.1492 | 0.1301 |
| SRO02_28133706 | 0.0732 | 0.1327 | 0.2356 | 0.2579 |
| SRO02_4535679 | 0.1697 | 0.1929 | 0.2084 | 0.1989 |
| SRO03_6110606 | 0.213 | 0.2463 | 0.1786 | 0.1358 |
| SRO04_14146782 | 0.0485 | 0.0517 | 0.1117 | 0.1719 |
| SRO05_13334757 | 0.3739 | 0.4313 | 0.3852 | 0.3477 |
| SRO06_24561982 | 0.3205 | 0.3486 | 0.3074 | 0.3013 |
| SRO07_9443780 | 0.3336 | 0.3225 | 0.2653 | 0.2165 |
| Total | 0.5135 | 0.585 | 0.5831 | 0.5826 |

**Table 3.** Cullis heritability and correlation with traditional traits for each spectral indices BLUPs with significant markers found in GWAS.

| Trait | Trait |  | Markers | Chromosome | Position | Minor Allele Frequency | P-value | Cullis Heritability | Correlation with traditional traits |  |  |  |
| --- | --- | --- | --- | --- | --- | --- | --- | --- | --- | --- | --- | --- |
|  | Method | Mask |  |  |  |  |  |  | Sev11 | Sev13 | Sev15 | Sev17 |
| Sev03 | median | NDVI | SRO01_3133661 | 1 | 3133661 | 0.4840426 | 3.72132E-07 | 0.00 | 0.11 | 0.11 | 0.08 | 0.09 |
| Sev03 | median | OSAVI | SRO01_3133661 | 1 | 3133661 | 0.4840426 | 3.72132E-07 | 0.00 | 0.11 | 0.11 | 0.08 | 0.09 |
| Sev07 | mean | NDVI | SRO04_30600585 | 4 | 30600585 | 0.4734043 | 1.366388E-07 | 0.12 | 0.23 | 0.2 | 0.16 | 0.18 |
|  |  |  | SRO07_6680558 | 7 | 6680558 | 0.4893617 |  |  |  |  |  |  |
| Sev07 | mean | OSAVI | SRO04_30600585 | 4 | 30600585 | 0.4734043 | 1.366388E-07 | 0.12 | 0.23 | 0.2 | 0.16 | 0.18 |
|  |  |  | SRO07_6680558 | 7 | 6680558 | 0.4893617 |  |  |  |  |  |  |
| Sev13 | mean | NDVI | SRO01_14234912 | 1 | 14234912 | 0.4893617 | 3.027627E-10 | 0.30 | 0.19 | 0.22 | 0.2 | 0.21 |
| Sev13 | mean | OSAVI | SRO01_14234912 | 1 | 14234912 | 0.4893617 | 3.027627E-10 | 0.30 | 0.19 | 0.22 | 0.2 | 0.21 |

### Resistance to late rust possibly has a locus of higher effect

The higher effect marker on 13 D.A.I., for both the network and the GWAS, had four candidate resistance genes flanking the marker. Resistance genes are involved in recognizing molecular patterns and elicitors of pathogens and are responsible for triggering the immune response in plants as soon as this recognition occurs. The “NB-ARC DOMAIN DISEASE RESISTANCE PROTEIN” gene, approximately 20kb from the marker, is a specific type of resistance gene that monitors intracellular disturbances related to plant immunity (van Ooijen *et al*. 2008). The “PLANT BROAD-SPECTRUM MILDEW RESISTANCE PROTEIN RPW8” gene was located at the exact location on the marker. RPW8 locus found in *Arabdopsis thaliana* contains a dominant resistance gene, which induces the salicylic acid metabolic pathway to trigger defense against fungal disease and powdery mildew (Xiao *et al*. 2001). The “WALLASSOCIATED RECEPTOR KINASE-LIKE 2” gene is 6.8kb from the marker. WAKs, or Plant cell wall-associated kinases, are genes in many plant species that bind to cell wall pectin. WAKs also have an intracellular domain for signal transduction and are related to the recognition of DAMPs, or Damage-Associated Molecular Patterns, as cell wall fragments. By recognizing wall integrity, these receptors can trigger callose deposition on the cell wall and increase protection against pathogens (Amsbury 2020). The only work studying late rust resistance in raspberry was the (Jamieson and Nickerson 1999) work, in which they experimented with segregating populations for late rust resistance. They concluded that there is a locus with a higher effect on resistance called “Pa” by them, agreeing with our results.

### The number of D.A.I. with higher resolution for disease severity

Although SRO05_13334757 SNP is significantly associated with resistance at 13 D.A.I., there are no significant markers at 11, 15 and 17 D.A.I. (Figure 5). This possibly happened because, besides being a small population size, the central parent (*R. occidentalis*) is not immune to *A. americanum*. Dias *et al*. (2023b) demonstrated that although the central parent has pre- and post-formed defense mechanisms against *A. americanum*, these are not sufficient to completely contain the infection. With the intense application of the inoculum to all the leaves and more than two weeks of disease progression, the disease severity scores were no longer representing the different resistance levels contained in the panel. Likewise, the disease had possibly not progressed far enough at 11 D.A.I. and the BLUPs did not represent the different resistance levels contained in the panel. Another indication that this occurred is that BLUPs averages are increasingly higher in time, and the standard deviation in 11 D.A.I. is smaller than those found in the other days (Figure 4). Furthermore, the highest heritabilities for disease severity found by the high-throughput phenotyping approach were at 13 D.A.I. With this, we conclude that the thirteenth D.A.I. is the highest resolution day for phenotyping of late rust resistance in raspberries.

### Gene flow throughout phenotyping days

An unexpected result was that, although both techniques have the SRO05_13334757 marker with higher association in the thirteenth D.A.I., only for GWAS does this same marker appear with a higher p-value in all traits. Meanwhile, the other phenotyping days in the network, Sev11, Sev15, and Sev17, are more correlated with other markers. Possibly, the relationship between graph separation and conditional independence between variables showed us these differences in the network, as we consider all variables (traits and SNPs) simultaneously, not as a univariate test as in GWAS. We also observed that the network could provide valuable insights into other genes with less effect on the evaluated trait. For example, we can mention the “probable WRKY transcription factor 2” gene on chromosome 6. Although this candidate gene is not a pathogen receptor, this transcription factor confers resistance to many diseases and abiotic stresses (Phukan *et al*. 2016). The “ankyrin repeat-containing protein NPR4” gene, on chromosome 7, is a receptor for salicylic acid, an important hormone in plant immunity (Liu *et al*. 2020). The “probable serine/threonine-protein kinase kinX” gene, on chromosome 3, is a constituent in the regulation of plant immunity signaling cascades (Ding *et al*. 2022), among many other genes found throughout the network that have functions directly linked to plant resistance.

### A guidance for raspberries breeding

Although our work has offered different confidence levels with the obtained results, it has produced important information for directing raspberry breeding programs. It was possible to indicate the marker with the highest effect in one of the main crop diseases using different statistical approaches; both led to consistent results, supporting our findings and conclusions. Furthermore, using a GWAS/copula graphical models pipeline, which is not conventional in plant breeding, can help other researchers discover causal alleles for their study traits. Using this pipeline, we could also add other candidate genes with less effect but still possibly responsible for the same trait. Beyond the traditional traits, we were able to indicate a new way and the highest resolution day for high-throughput phenotype raspberries. Therefore, a natural progression for this work would be to carry out validation studies of the markers found to confer reproducibility and consistency, as no other works are studying the genetic control of late leaf rust in raspberries to compare significant markers.

## Datax Availability

Phenotypes are available upon request. Genomic data in Variant Call Format (VCF) is available in the Rosaceae database, under the accession number tfGDR1071, under the name “Rocci_raw_imput_all_sort.vcf” and can be accessed with the URL https://www.rosaceae.org/publication_datasets.

## Acknowledgments

We would like to express our acknowledgement to the *Fundação de Amparo à Pesquisa do Estado de São Paulo* (FAPESP), for the financial support provided through the 19/13191-5 thematic project.

